# Functional diversification of UBP6 in plant immunity through N-degron pathway regulation

**DOI:** 10.64898/2026.07.24.740541

**Authors:** Charlene Dambire, Gabriel Sánchez, Yovanny Izquierdo, José R. Valverde, Isabel Manrique, Álvaro D. Fernández-Fernández, Kris Gevaert, Frank Van Breusegem, Neil J. Oldham, Michael J. Holdsworth, Jorge Vicente

**Affiliations:** School of Biosciences, University of Nottingham, LE12 5RD, UK; School of Chemistry, University of Nottingham, University Park, Nottingham NG7 2RD, UK; Department of Plant Molecular Genetics, Centro Nacional de Biotecnología CSIC, Campus of Cantoblanco, 28049 Madrid, Spain; Department of Plant and Microbial Biology, University of Zürich, 8008 Zürich, Switzerland; Department of Biomolecular Medicine, Ghent University, Technologiepark 75, 9052, Ghent, Belgium, VIB-UGent Center for Medical Biotechnology, VIB, Technologiepark 75, 9052, Ghent, Belgium; Department of Plant Biotechnology and Bioinformatics, Ghent University, 9052, Ghent, Belgium, Center for Plant Systems Biology, VIB, Technologiepark 71, 9052, Ghent, Belgium

## Abstract

Deubiquitylases are key proteolytic regulators of ubiquitin-dependent cellular processes, catalyzing the removal or remodelling of ubiquitin modifications on substrate proteins, including those targeted for proteasomal degradation. UBIQUITIN PROTEASE (UBP)6 is a deubiquitylase that promotes the abundance of NONEXPRESSOR OF PATHOGENESIS RELATED GENES (NPR)1, a conserved master regulator of plant immunity. Here, we show that the *Arabidopsis thaliana* protease METACASPASE (MC)9 site-specifically processes UBP6, generating the E157-UBP6 proteoform, whose stability is controlled by the Arginyl-transferase (ATE) N-degron pathway. We observed that pathogen recognition both triggers UBP6 cleavage and leads to conditional stabilisation of E157 UBP6, which is enhanced as the defence response intensifies. Our data suggest that E157 UBP6, which lacks deubiquitylating activity, may induce inhibition of the proteasome, elevating NPR1 levels and enhancing salicylic acid (SA) induced gene activation, all of which collectively contribute to restricting pathogen growth. Thus, UBP6 cleavage and N-degron pathway regulation provide distinct proteoforms of UBP6 with specific effects on the immune processes.

## Introduction

Plants constantly interact with pathogenic organisms, establishing coordinated responses that operate at multiple levels and that are subject to constant evolution^1^. Plants must be able to deploy a specific protein portfolio tailored to each pathogenic challenge. However, protein homeostasis (proteostasis) undergoes significant imbalances during pathogenic attack^2^. Control of the stability of plant defence proteins has been identified as an important regulator of their activity, is crucial for the response against pathogens and, therefore, is a target for pathogenic machineries ^3–5^.

The ubiquitin-proteasome system or UPS is a major regulator of protein half-life, acting through the attachment of ubiquitin to substrates to alter their function or mark them for degradation by the 26S proteasome complex^6,7^. Within the UPS, the N-degron pathways, form a substrate-recognition branch that enhance protein degradation by recognition of an amino-terminal (Nt-) destabilizing residue^8,9^. Such residues are exposed by peptidase action and recognized by a set of Nt-modifying enzymes and E3 ligases. N-degron pathways in plants control multiple abiotic and biotic stresses, for example the NTAQ1, PCO and PRT1 N-degron pathways modulate different processes of the immune response^10–15^ (Fig. S1a).

Destabilising Nt-residues are produced by protease cleavage^16^, producing Nt- and carboxy-terminal (Ct-) proteoforms^17^. Therefore, the combined activities of peptidases and N-degron targeting can determine protein abundance and activity. Remarkably few *in vivo* substrates of N-degron pathways are known in any system^18–20^, primarily because the relevant protease–substrate relationships are not known. Metacaspases are plant caspase-like proteases with proven roles in plant defence^21–25^. It was reported that Arabidopsis METACASPASE (MC)9 cleaves more than 380 proteins^17,26^. Here we show that one of the proteoforms resulting from MC9 cleavage, Glu(E)157-UBIQUITIN PROTEASE (UBP)6, is a substrate of the ATE N-degron pathway. UBP6 is a deubiquitylase that regulates the abundance of the master regulator of plant defence NONEXPRESSOR OF PATHOGENESIS RELATED GENES (NPR)1^27^. Although E157-UBP6 lacks deubiquitylating activity, it retains proteasome-interacting capacity and acquires a distinct non-catalytic function. Upon immune activation, UBP6 cleavage is induced and E157-UBP6 becomes conditionally stabilized, leading to NPR1 accumulation, enhanced salicylic acid-responsive gene expression, and increased resistance to *Pseudomonas syringae*. These findings reveal proteolytic processing and N-degron regulation as mechanisms that diversify UBP6 function, allowing a conditionally stable proteoform to modulate proteasome activity and reinforce plant defence.

## Results

### The ATE N-degron pathway regulates the half-live of the MC9-derived proteoform E157-UBP6

A previously published N-terminome dataset showed that MC9 cleaves UBP6 at position R156, generating the Ct-proteoform E157-UBP6^17^. Glutamic acid (E) is a destabilizing residue that, as part of an N-degron, can be arginylated by ATEs, making E157-UBP6 a potential substrate of the ATE N-degron pathway^8^ (Fig. 1a). We confirmed in two orthogonal assays that UBP6 is an MC9 target and subsequently analysed whether the generated proteoform E157-UBP6 is a substrate of the ATE N-degron pathway. *In vitro* incubation in a rabbit reticulocyte lysate (RRL) of recombinant UBP6 in the presence of recombinant MC9 and the proteasome inhibitor bortezomib (BZ) produced a band of the expected size for E157-UBP6-3HA (Fig. 1b). *In vivo* incubation of UBP6 with recombinant Wild Type (WT) MC9 in *Nicotiana benthamiana* produced a band of the expected size for E157-UBP6, while incubation with the catalytically inactive mutant MC9^CACA^^28^ did not (Fig. 1c). An *in vitro* analysis in a RRL using the Ubiquitin Fusion Technique (UFT)^29^ construct UFT-E157-UBP6-3HA (Fig. S1b), hereafter E157-UBP6, demonstrated a shorter half-life for this carboxy-terminal (Ct-) proteoform, that was repressed by BZ (Fig. 1d). *In vitro* coupled transcription/translation of E157-UBP6-3HA in a wheat germ extract (that does not contain an active proteasome) followed by mass spectrometry analysis showed that the fragment is N-terminally arginylated (Fig. 1e).

**Figure 1:**
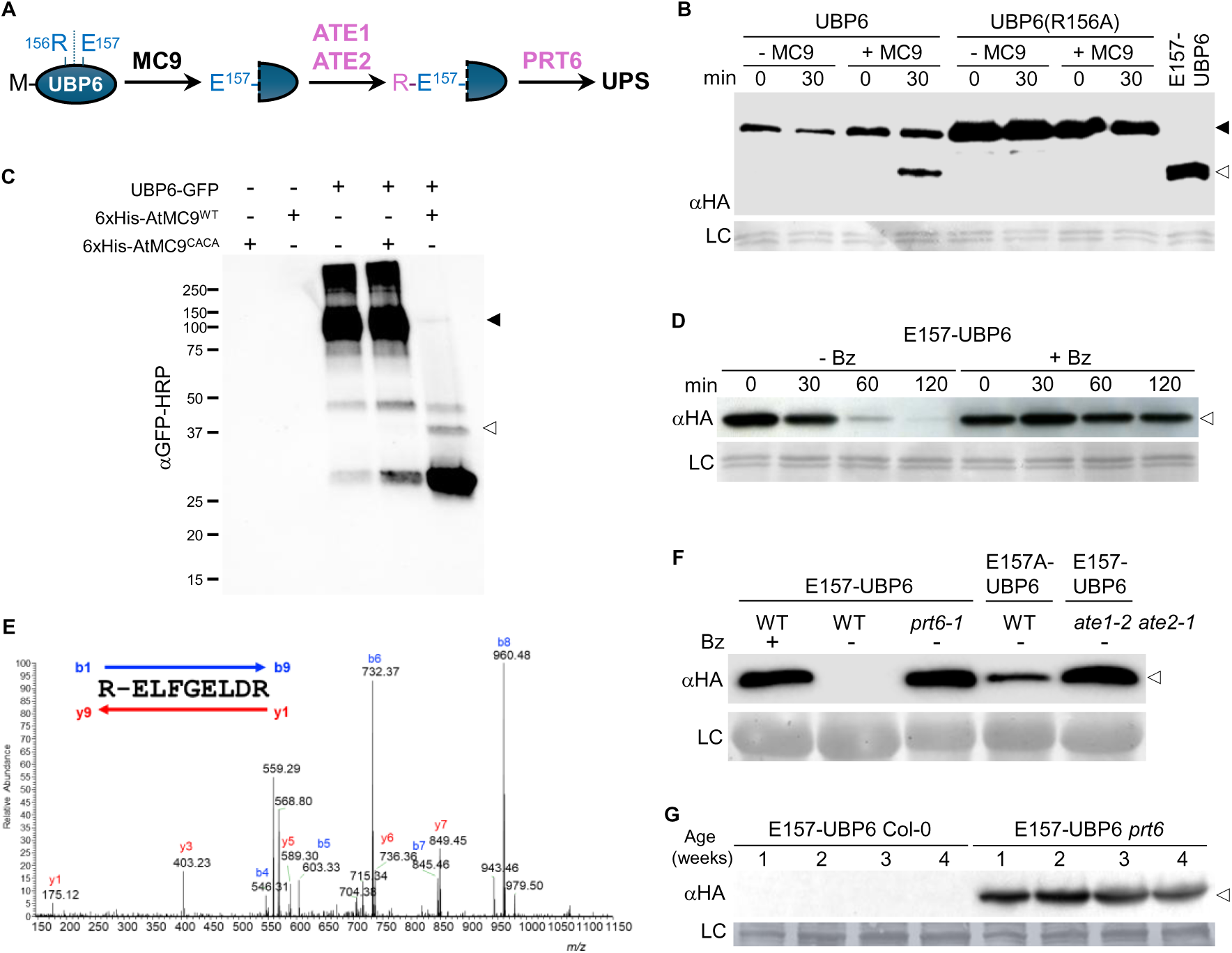
UBP6 is a substrate of both METACASPASE9 and the ATE N-degron pathway. (A) Schematic presentation of the UBP6 processing by MC9 and ATE N-degron pathway. Single amino acid codes are used; ATE, arginyl-transferase; PRT6, PROTEOLYSIS6; UPS, Ubiquitin Proteasome System. E157-UBP6, substrate of MC9 and the ATE N-degron pathway. (B) *In vitro* cleavage of UBP6 by MC9. Comparison of time course of cleavage of UBP6 with UBP6(R156A) that abolishes the MC9 P1 recognition site. (C) UBP6 purification from *Nicotiana benthamiana* showing specific proteolysis in presence of active MC9. (D) Analysis of E157-UBP6 stability over time in rabbit reticulocyte lysate in the presence or absence of bortezomib (Bz). (E) ESI-MS/MS spectrum assay showing arginylation of the proteoform E157-UBP6 transcribed and translated using a wheat germ extract. (F) *In vivo* analysis of E157-UBP6 stability in WT and N-degron pathway *prt6-1* and *ate1-2 ate2-1* mutant seedlings. (G) Comparison of stability of E157-UBP6 in WT and *prt6-1* plants during different developmental stages. Ponceau staining of Western blots is shown (LC: loading control). Closed triangle, full length UBP6; Open triangle, HA-tagged E157-UBP6 Ct-proteoform.

We analysed the N-degron pathway dependent-stability of E157-UBP6-3HA *in vivo* in transgenic Arabidopsis. The proteoform was barely detectable in WT seedlings but was stabilised in the presence of bortezomib (Fig. 1f). We observed stabilization of E157-UBP6 in both *ate1 ate2* and *prt6-1* mutant seedlings and of E157A-UBP6, where the Nt-E was replaced by Ala (A), a non-destabilising residue. The proteoform was also constitutively stabilised in a *prt6-*1 background during 4 weeks of plant growth (Fig. 1g). Together, these data show that UBP6 is cleaved by MC9 and the resulting fragment E157-UBP6 is a substrate of the ATE N-degron pathway (Fig. 1a).

### *Pseudomonas* detection induces the cleavage and conditional stabilization of E157-UBP6

In unchallenged conditions, NPR1, a known target of UBP6^27^, localizes in the cytoplasm as inactive oligomers. A surge in salicylic acid (SA) levels following pathogen recognition leads to a cellular redox change that induces partial NPR1 monomerization and translocation to the nucleus where it serves as a coactivator of the expression of SA-dependent defence genes^30^. We observed that treatment with SA strongly increased the stabilization of E157-UBP6 in *ubp6 ubp7* (hereafter *u6u7*) mutant seedlings (UBP7 is a paralog of UBP6, so it is necessary to use the double mutant to achieve total function removal) (Fig. 2a). The mutated proteoform E157R-UBP6, with Nt-R (changing substrate specificity from ATE to PRT6; Fig. 1a), showed the same instability in *u6u7* as E157-UBP6, with stability increasing after SA treatment. As expected, E157-UBP6 showed increased stabilization in the triple mutant *prt6-1 ubp6-1 ubp7-1* (hereafter *p6u6u7*), where the proteoform cannot be degraded through the N-degron pathway.

**Figure 2.**
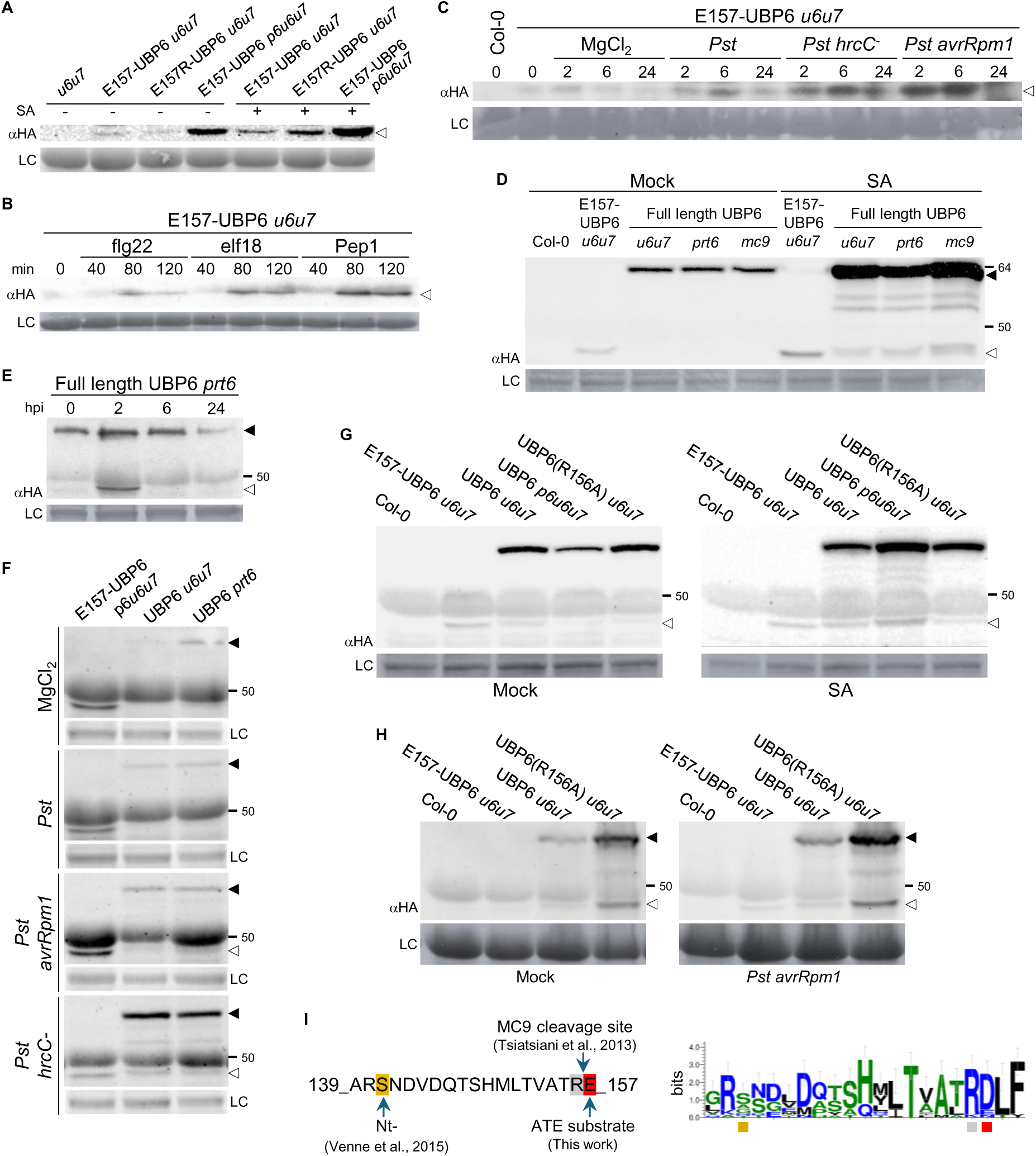
Conditional stability of E157-UBP6 during activation of the immune response. (A) WB showing E157-UBP6 accumulation by salicylic acid (SA) and that shifting ATE dependency of E157-UBP6 to PRT6, E157R-UBP6, results in an equally unstable proteoform. (B) E157-UBP6 transiently accumulates in Arabidopsis *u6u7* mutant seedlings following treatment with P/DAMPs. (C) E157-UBP6 transiently accumulates in Arabidopsis *u6u7* mutant plants following inoculation of *Pst* strains that elicit different levels of resistance. (D) Endopeptidase cleavage of full length UBP6 and stabilization of the resulting proteoform E157-UBP6 in seedlings is activated by SA and (E) inoculation of virulent *Pst* in adult plants. (F) Stability of UBP6-related proteoforms at 24h after inoculating *Pst* strains. (G) Stability of UBP6-related proteoforms in UBP6(R156A) line in SA-treated seedlings and (H) inoculated with *Pst avrRpm1* adult plants. (I) Schematic presentation of the UBP6 protein sequence subjected to several N-terminal modifications (left) and the degree of conservation of each amino acid (right). Closed triangle, full length UBP6; Open triangle, HA-tagged E157-UBP6 Ct-proteoform. Ponceau staining of Western blots is shown (LC: loading control).

During a pathogenic attack, one of the earliest activated processes is the recognition of conserved pathogen structures (PAMPs) by membrane receptors, which activates pattern triggered immunity (PTI), a tightly regulated response in which the N-degron pathways are involved^13^. Stabilization of E157 UBP6 increased in response to the PAMPs flg22 and elf28 and the damage associated (DAMP) Pep1 (Fig. 2b). In line with this, we observed increased expression of PAMP responsive genes in response to flg22, though not to elf28 or Pep1, in *prt6* mutants (Fig. S2a,b). Expression of the N-degron pathway genes *PRT6*, and *ATE1* in WT seedlings showed only a slight increase following flg22, elf18 and Pep1 treatment (Fig. S2c).

We next investigated whether immunity in a broader context, particularly during infection by the virulent bacterial strain *Pseudomonas syringae* pv. *tomato* DC3000 (hereafter *Pst*), impacts the stability of E157 UBP6. Recognition of *Pst* by the plant induced transient stabilization of E157 UBP6 (Fig. 2c). We observed stronger transient stabilization of the proteoform during infections with the mutant *Pst hrcC* and the avirulent *Pst avrRpm1* strains compared with the virulent *Pst*. *Pst hrcC* is defective in the Type III Secretion System (T3SS), preventing injection of virulence effectors into host cells and therefore acting essentially as a PTI only inducer. *Pst avrRpm1* expresses the *avrRpm1* gene, which is recognized by the Arabidopsis RPM1 R gene and triggers a rapid and strong defence response (effector triggered immunity; ETI), characterized by a hypersensitive response (HR). Analysis of *MC9* gene expression levels in seedlings treated with SA showed a decrease (Fig. S2d), while little or no changes were observed in plants inoculated with *Pst*, *Pst hrcC-* and *Pst avrRpm1* bacterial strains, respectively (Fig. S2e). Together, these results indicate a positive correlation between the strength of the immune response and the degree of stabilization of E157 UBP6.

To determine the relationship between MC9 activity and E157-UBP6 formation, we analysed processing of UBP6 *in vivo* by expression of the construct UFT-UBP6-3xHA. Cleavage of UBP6 was triggered by exogenous application of SA in *u6u7* and *prt6-1* plants, generating a fragment of the same size as E157A-UBP6-3HA (Fig. 2d). Cleavage was not abolished in *mc9* mutants, suggesting redundancy with other metacaspases that may also cleave UBP6.

In addition to E157-UBP6-3xHA a slightly higher molecular weight band was also observed, that might indicate a closely associated upstream cleavage site in UBP6, although no other cleavage sites in UBP6 were found in the MC9 N-terminome dataset. Importantly, inoculation of *Pst* in adult plants also triggered the cleavage of UBP6 (Fig. 2e). In line with the previously described increased accumulation of E157-UBP6 during infections with *Pst hrcC* and *Pst avrRpm1* (Fig. 2c), inoculation of these pathogens also induced the cleavage of UBP6, but the resulting proteoform remained stable at 24 h, whereas it was already degraded in infections with virulent *Pst* (Fig. 2f).

To further understand the impact of E157 UBP6 generation and stabilization on plant defence, we analysed plants unable to produce this proteoform by transforming *u6u7* with a full-length UBP6 in which the MC9 cleavage site R156 was mutated to Alanine (UBP6(R156A)). Surprisingly, UBP6(R156A) showed a proteoform with a similar size than E157-UBP6 in both seedlings treated with SA (Fig. 2g) and adult plants inoculated with the avirulent strain *Pst avrRpm1* (Fig. 2h). This suggests that an alternative cleavage site near the residue R156 may exist in UBP6, or that another protease is able to cleave in the presence of A156.

Previous reports have highlighted the importance of the region spanning residues A139 to R156 for UBP6 regulation^31^. In addition to the MC9 cleavage site at R156^17^, S141-UBP6 was identified in a proteomic screen for protease targets^31^(Fig. 2i). In this context, R140 may also represent a metacaspase cleavage P1 site. Residues associated with cleavage at positions 140 and 156, as well as the destabilising residue at 157, are conserved across plant and non plant UBP6 homologues (Fig. 2i, S2f). In yeast the structure of Ubp6 bound to the proteasome has recently been described^32^. Our analysis shows that yeast and Arabidopsis UBP6 structures in that area are highly similar (Fig. S2g). Structural prediction reveals that the S141–R156 region forms a loop followed by an α-helix. Together, these observations suggest that the proteoform detected in UBP6(R156A) might correspond to S141 UBP6 3HA (∼41.8 kDa), rather than E157 UBP6 3HA (∼40.1 kDa). This proteoform may serve as stable precursor of the active E157 UBP6 generated during pathogen recognition, or alternatively, loss of cleavage at R156A may activate other sites not normally used.

### E157-UBP6 participates in the immune response against *Pst*

MC9 cleavage of UBP6 at R156 separates the deubiquitylase active site at C113 from the Ct-proteoform. Interestingly, it was already described that UBP6 in Arabidopsis and yeast also has a non-catalytic function based on its ability to directly interact with the proteasome and allosterically inhibit its activity^27,32,33^. To understand the impact that cleavage of UBP6 and stabilization and function of E157-UBP6 have on the immune response, we examined, in the *prt6* background, the interactome of E157 UBP6 compared with that of full-length UBP6 in the presence or absence of SA. Immunoprecipitation of the bait proteins expressed in seedlings followed by mass spectrometry analysis indicated both overlapping and unique interactions among constructs and treatments (Fig. 3a, S3a; Data S1). Functional analysis showed a cluster of interactors composed of several proteasome subunits (Fig. 3b), among which RPN6, already described as an interactor of UBP6^27^(Fig. S3b,c). We observed an increased number of proteasome subunits associated with UBP6 after SA treatment, whereas in the case of E157 UBP6, the cluster contained fewer subunits and remained almost unchanged between treatments. In mock conditions the presence of the E157-UBP6 fragment significantly increased ubiquitylation levels (Fig. 3c,d, S3d), reinforcing the idea that a capacity to interact with proteasome subunits may exert an inhibitory effect on proteasome function. As previously reported, ubiquitylation increased in WT plants after SA treatment^34^ (Fig. S3d). Unlike E157-UBP6 *u6u7*, UBP6 *u6u7* and UBP6(R156A) *u6u7* did not show increased total ubiquitylation in untreated seedlings compared with *u6u7* (Fig. 3d, S3d). After SA treatment or inoculation with the avirulent strain *Pst avrRpm1* (Fig. 3d, S3d), a similar general increase in ubiquitylation was observed in every line, suggesting that the effect observed in the untreated E157-UBP6 *u6u7* is masked by the activation of additional proteasome regulatory mechanisms. Together, this data reinforces the idea that E157-UBP6 is an active fragment generated during the immune response, where both full length and proteoform may exist developing complementary activities.

**Figure 3.**
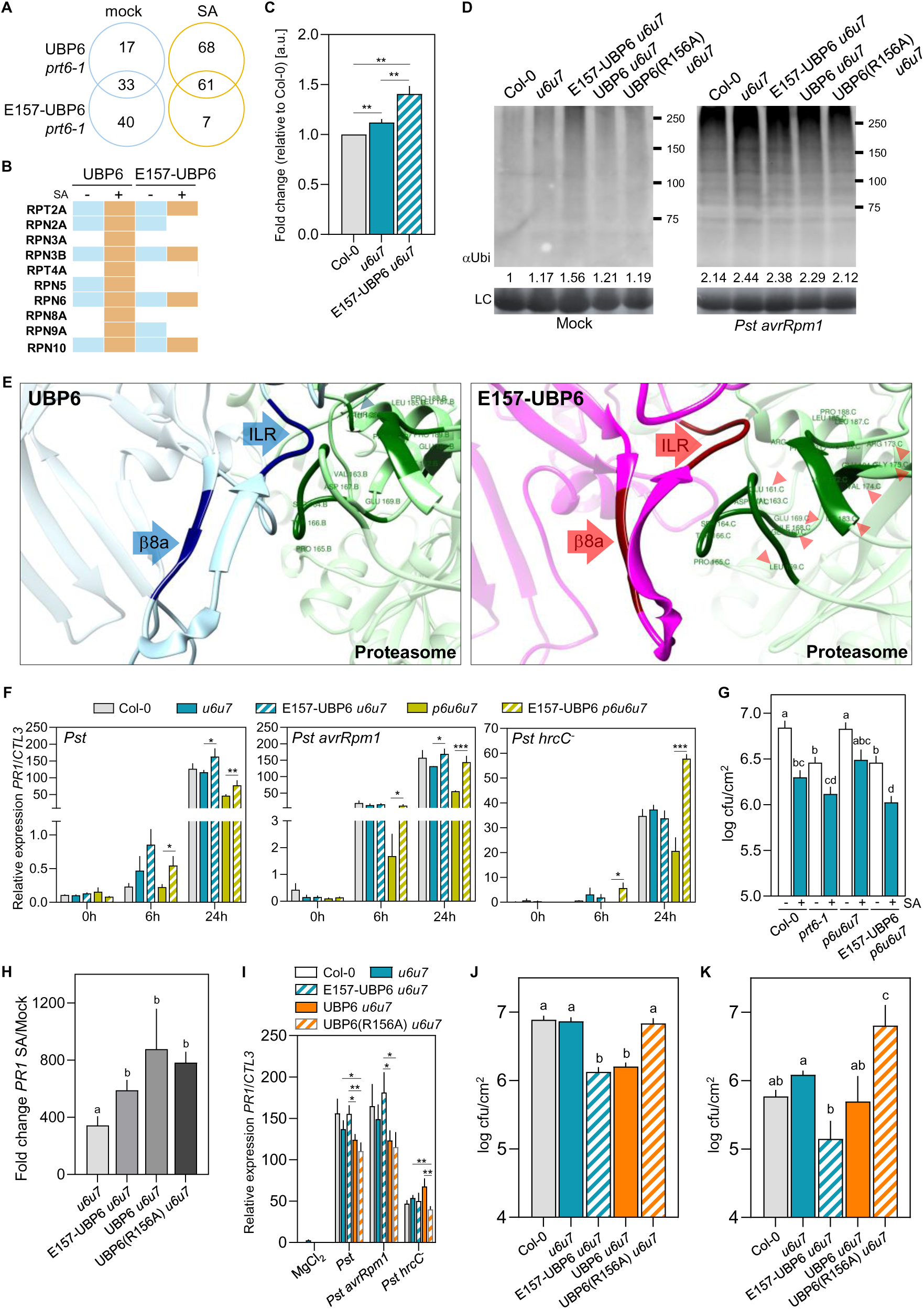
E157-UBP6 participates in the immune response against *Pst*. (A) Proteins found to associate with proteoform E157-UBP6 and full length UBP6 following mock (blue) or SA (orange) treatment in Co-IP assays. (B) Interaction dynamics of E157 UBP6 and UBP6 with 26S proteasome subunits following mock (blue) or SA (orange) treatment. (C) Quantification of total protein ubiquitylation observed in immunoblots with anti-Ub antibody in untreated seedlings. (D) Immunoblot with anti-Ub antibody showing total ubiquitinated proteins in adult plants inoculated with *Pst avrRpm1*. (E) Detail of the structure of UBP6 (light blue) and of E157-UBP6 (pink) bound to the proteasome (green). β8a and ILR motifs are coloured in a darker shade and their location is indicated by labelled arrows. The a.a. in the proteasome that interact with them are indicated in a darker green shade. Non-labelled arrows indicate a.a. that interact differentially between UBP6 (dark blue) and E157-UBP6 (red). (F) qRT-PCR analysis of NPR1-dependent gene *PR1* expression in plants inoculated with *Pst* strains. (G) Quantification of *Pst* growth 72 h after bacterial infiltration in plants pretreated with H_2_O or 0.5 mM SA 24 h earlier. (H) qRT-PCR analysis of *PR1* gene expression in seedlings treated with SA and (I) inoculated with *Pst* strains. (J) Quantification of *Pst* growth 72 h after bacterial infiltration in adult plants. (K) Quantification of *Pst* growth 72 h after bacterial surface-spray in adult plants. Statistical differences were analysed by ANOVA followed by Tukey test (P < 0.05), significant differences are indicated with letters, or Student’s t-test: *, P < 0.05; **, P < 0.01; ***, P < 0.001. Ponceau staining of Western blots is shown (LC: loading control).

Yeast ScUbp6 binding to the proteasome and ubiquitin is mediated by its BL1 loop, a specific sequence that contains three elements, the ILR motif and the β8a and β8b strands, that induces allosteric inhibition of the proteasome^32^. This interaction triggers a series of rearrangements in downstream structural elements, namely BL2 and SL, ultimately leading to proteasome inhibition. The ILR element is highly conserved in plants (Fig. S3e). Using the publicly available Ubp6–proteasome structure^35^, our analysis shows that the BL1 structures of yeast and Arabidopsis UBP6 are highly similar and that the MC9 cleavage site is exposed (Fig. S3f,g). Comparison of Arabidopsis E157-UBP6 with full-length UBP6 revealed differences in the folding of the ILR element (the proteasomal contact region) and in the β8a strand, the element that occludes ubiquitin (Fig. 3e, S3h). Our analysis showed that E157-UBP6 interacts with a larger area of the proteasome, specifically with 7 more amino acids of the subunits RPN6 and RPT1, than UBP6. This enhanced interaction may induce a stronger conformational change in the proteasome, leading to increased inhibition and, consequently, higher levels of cellular ubiquitylation (Fig. 3c,d, S3d).

Stabilization of E157 UBP6 in *p6u6u7* during the immune response resulted in increased expression of the SA responsive defence marker *PR1* (Fig. 3f) and extended activation of the MAPKs MPK3/6 (Fig. S3i). We next analysed the impact of E157-UBP6 in the broader context of pathogenic infection by *Pst*. This showed that plants expressing E157 UBP6 had enhanced resistance to *Pst*, an effect reinforced in SA pre-treated plants (Fig. 3g).

Although *PR1* expression increased in non-cleavable UBP6(R156A) *u6u7* seedlings to levels similar to those observed in E157 UBP6 *u6u7* and UBP6 *u6u7* after SA treatment (Fig. 3h), it failed to reach comparable levels in adult plants inoculated with the three bacterial strains differing in immunity induction strength (Fig. 3i). UBP6(R156A) *u6u7* failed to restrict the growth of injected virulent *Pst* to the same extent as native UBP6 (Fig. 3j) and showed even higher bacterial levels when the pathogen was sprayed onto the leaf surface (Fig. 3k). This implies that cleavage of UBP6 and stabilization of E157-UBP6 constitute an important component of UBP6 action.

### Conditional stability of E157-UBP6 affects the expression of NPR1-dependent and independent genes

We analysed whether E157-UBP6 retains control over NPR1 action in immunity. As previously described^27^, *u6u7* showed a reduction in NPR1 accumulation after SA treatment (Fig. 4a), whereas in E157-UBP6 *u6u7* we found NPR1 levels to increase. It has been described that UBP6, apart from its deubiquitylating activity, induces a non-catalytic proteasome inhibition promoting the expression of a specific subset of NPR1-dependent genes^27^, specifically *PR* genes. E157-UBP6 in both *u6u7* and *p6u6u7* showed increased expression of several *PRs* and *WRKYs* compared with the non-transgene multiple mutants (Fig. 4b). Given the positive role of the E157 UBP6 proteoform in the accumulation of NPR1 (Fig. 4a), we analysed the impact of stabilization of this proteoform on the transcriptome in response to SA. We performed an RNA-seq analysis after treatment with SA of E157-UBP6 in *p6u6u7* compared with *p6u6u7*. We observed a significant overlap between the two lines of both SA-induced and repressed genes, including a significant percentage of NPR1-dependent genes^27^, compared with their respective mock-treated samples (Fig. 4c; Data S2), Regarding the shared NPR1-dependent genes (255 induced and 91 repressed), we observed differential expression changes associated with the presence of stable E157 UBP6 (Fig. 4d), reinforcing its role in regulating NPR1 stability and transcriptional coactivator activity. For both NPR1-dependent and independent genes induced by SA, the fold changes between the two lines were also compared, showing that in both cases induction was significantly stronger in the presence of stabilised E157-UBP6 (NPR1-dependent p-value 1.263e-10; NPR1-independent p-value 5.012e-12) (Fig. 4e), and also observed that the induction caused by the proteoform was significantly higher in the NPR1-dependent genes compared with NPR1-independent genes (p-value 0.00076). Together, these data show that E157-UBP6 causes a consistent general increase in SA-responsive gene induction, which is pronounced in NPR1-dependent genes.

**Figure 4.**
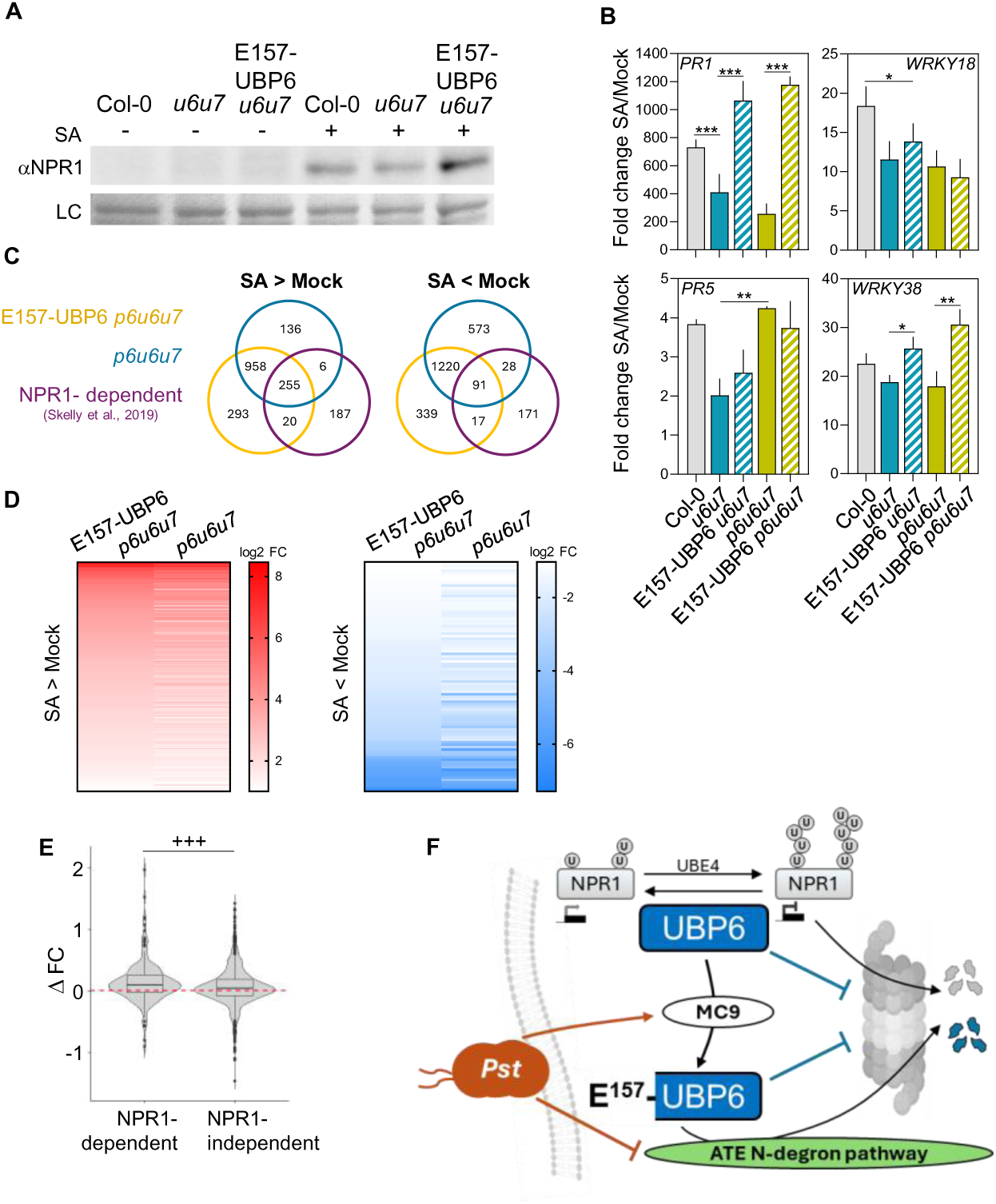
E157-UBP6 adjusts NPR1-dependent and -independent transcriptome. (A) NPR1 protein levels in Arabidopsis seedlings after treatment with SA. (B) qRT-PCR analysis of NPR1-dependent genes *PR1* and *WRKY18* expression in seedlings treated with SA. Statistical differences were analysed by Student’s t-test: *, P < 0.05; **, P < 0.01; ***, P < 0.001. (C) Venn diagrams showing differentially expressed genes in the indicated lines mock or SA-treated. (D) Comparison between E157-UBP6 *p6u6u7* and *p6u6u7* lines of the gene expression values of NPR1-dependent genes up (red) or down (blue) regulated after SA treatment. (E) Distribution of delta increase between the fold changes of NPR1-dependent and independent genes in E157-UBP6 *p6u6u7* vs *p6u6u7* (ΔFC= E157-UBP6 *p6u6u7* - *p6u6u7*). Within each gene set, statistical differences were analysed by paired Wilcoxon test. Between both gene sets, statistical differences were analysed by non-paired Wilcoxon test: +++, P < 0.001. (F) Model showing generation and function of E157-UBP6 during plant defence against *Pst*: (1): Pathogen recognition triggers salicylic acid accumulation and MC9 activation; (2): MC9 cleaves UBP6, generating the proteoform E157-UBP6. The ATE N-degron pathway recognises and degrades it to prevent an unnecessary immune response; (3): As part of the activated defence state, immune signals temporarily inhibit the ATE N-degron pathway, E157-UBP6 becomes stable and binds to the 26S proteasome, significantly reducing its proteolytic activity; (4): Reduced proteasome activity leads to the accumulation of NPR1 and other defence proteins, triggering the expression of defence genes and enhancing resistance; (5): Once the infection subsides, the ATE N-degron pathway becomes active again, degrading E157-UBP6 to restore normal proteasome activity and basal immune levels. Ponceau staining of Western blots is shown (LC: loading control).

## Discussion

It was previously shown that UBP6 has two distinct activities: a catalytic activity, deubiquitylation, and a non catalytic one, allosteric inhibition of the proteasome^32,27^. Both activities lead to inhibition of the degradation of the master transcriptional co-activator NPR1, but surprisingly they result in different outcomes, including the activation of distinct subsets of NPR1-dependent genes^27^. How does the cell initiate these two activities? Our data suggests that this occurs through the formation of the E157-UBP6 proteoform. The surge of SA levels upon pathogen infection produces a redox cellular change resulting in a decrease of pH, to between 5.8 and 5.4^36^. Interestingly, MC9 becomes active at acidic pH, between 5.6 and 5.0^37^. We suggest the following temporal mechanism of UBP6 action. Upon pathogen recognition, UBP6 deubiquitylates NPR1 thereby maintaining SA signalling in an active state. At a later stage of infection, probably when the cytoplasm reaches a low enough pH due to cellular redox change, MC9 (and perhaps other MCs^38^) are activated and UBP6 is cleaved, generating E157 UBP6. The cleavage and stabilization of the fragment is detectable at two hours of infection, sooner in the case of treatment with flg22, which indicates that cleavage can occur rapidly and, subsequently, it is the inhibition of the ATE N-degron pathway that determines proteoform accumulation. Results obtained comparing E157-UBP6 and E157R-UBP6 indicates that inhibition of the pathway occurs downstream of ATE (Fig. 1a) perhaps through repression of PRT6 activity. E157 UBP6 levels fluctuate over the course of infection, indicating that changes in the activity of the N-degron pathway are dynamic. Cell extracts containing the catalytically inactive E157 UBP6 in *u6u7* displays higher levels of total ubiquitination than *u6u7* under basal conditions. On the other hand, full length UBP6 *u6u7* shows slightly higher levels than non-cleavable UBP6(R156A) *u6u7* in response to SA treatment or bacterial infection. Once E157-UBP6 is generated and conditionally stabilised, our data shows that it binds to the proteasome. An *in silico* analysis indicated that this interaction is stronger with E157-UBP6 than with native UBP6, which could lead to a more robust, but still reversible, allosteric inhibition. Later during infection E157 UBP6 is degraded through the proteasome. This relieves proteasome inhibition, allowing the degradation of NPR1, and other potential UPS substrates. Our data indicates that the generation and stabilization of E157-UBP6 provide a mechanism to indirectly maintain activation of a subset of NPR1 dependent genes while simultaneously promoting the activation of distinct NPR1 independent gene sets.

Our study thus suggests that E157 UBP6 is a functional proteoform generated and controlled by protease and N-degron pathway activities during the plant defence response to provide two UBP6 functions, amplifying SA dependent defence, leading to enhanced resistance against *Pst*. This aligns with recent work showing that basal levels of SA repress the PRT6 N-degron pathway to coordinate growth and responses to abiotic stress in rice^39^. In a broader context, E157-UBP6 stabilization may result in the accumulation of immune-related proteins, not just NPR1, that in basal conditions are rapidly degraded under non-challenged conditions to prevent unnecessary defence activation. E157-UBP6 directed-proteasome inhibition may also interfere with pathogen strategies that hijack ubiquitylation to degrade host defence proteins. Increased E157 UBP6 stability during *Pst hrcC-* infections hints that pathogenic effectors may target components of the N-degron pathways to prevent substrate stabilization. The proteoform remained stable for longer during both *Pst hrcC-* and *Pst avrRpm1* infections, suggesting that it may contribute to the increased immune response that restricts the growth of these strains and/or that these pathogens are unable to re induce its degradation. It will be interesting in the future to analyse whether E157-UBP6 is also involved in proteasome inhibition leading to the accumulation of misfolded or damaged proteins, which could act as a stress signal, or in unbalancing hormone pathways regulated by UPS, thereby inducing a shift toward defence activation.

Our analyses have shown that key residues involved in cleavage of UBP6 and stabilization of E157-UBP6 are highly conserved across plant and non plant UBP6 orthologues. This is particularly relevant as the N-degron pathways are conserved in plant and animals and the human UBP6 orthologue USP14, a protein with an important role in cancer and neurodegenerative diseases such as Parkinson and Alzheimer, has conserved MC9 P1(R)-P1’(D) sites (Fig. S2f). This suggests a potential role of enzymes involved in neurodegenerative diseases that cleave after R, as Kallikrein-Related Peptidases (KLKs). This study might therefore help to elucidate USP14 role in maintaining the balance between the pool of free ubiquitin (catalytic function) and proteasome inhibition (noncatalytic function).

## Methods

### Plant materials and growth conditions

All work was carried out in the *A. thaliana* Col-0 accession background. Double mutant *ubp6 ubp7* was kindly donated by Dr. Steven Spoel (University of Edinburgh). Plants were grown and assays performed in controlled-environment rooms under the following conditions: 14 h of light (22 °C) and 10 h of dark (18 °C), 60–70% relative humidity. Plants were treated between 3 and 4 weeks after germination. For the analysis of Arabidopsis response *in vitro*, seeds were sowed on solid half-strength MS and, after 8 days, were transferred to 6-well tissue culture plates containing 4 ml of half-strength MS and left 24 h on a flatbed shaker (60 rpm) to acclimate. Then seedlings were treated with bortezomib 25 μM (Santa Cruz Biotechnology), SA 0.5 mM (Sigma-Aldrich), flg22 1 μM (GenScript), elf18 1 μM (Biomatik), Pep1 1 μM (Biomatik), and material was collected at specific times indicated in the text.

### Generation of ubiquitin fusion technique constructs

cDNA was synthesised from RNA extracted from 7-day old *Arabidopsis thaliana* seedlings and used as a template to PCR amplify UBP6 starting at E157 and cloned into a FLAG-DHFR-UBIQUITIN-3HA vector as previously described^40^. As described in ^41^ this provides a SacII restriction site downstream of the UBIQUITIN moiety that allowed cloning in frame of test proteins with defined amino-terminal residues. In this case we cloned UBP6 starting at E157, so that upon cleavage *in vivo* by constitutive deubiquitinating enzymes E157-UBP6-3HA or similar proteoforms (Extended data Fig. 2) are released from the pre-protein. This construct was subcloned into plant transformation gateway vector pH7wg2^42^ providing constitutive expression from the 35S CaMV promoter. Plants were transformed as previously described^43^.

### Arginylation of E157-UBP6

Arginylation of the proteoform E157-UBP6 was analysed by *in vitro* affinity purification, SDS-PAGE and gel band excision, in-gel protein digestion and mass spectrometry analysis of proteoform digests as previously described^40^.

### Site-directed mutagenesis

Site-directed mutagenesis primers were designed using the QuikChange Primer Design webpage and reactions performed using the protocol based on DpnI digestion of methylated DNA^44^. Positive clones were identified by colony PCR and confirmed by DNA sequencing. All primers used are listed in Extended Data Table 1.

### *In vitro* protein stability analysis

The stability of proteins was assessed *in vitro* using the rabbit reticulocyte lysate (RRL) coupled transcription/translation system as previously described, this heterologous system has previously been shown to contain all components of the Arg/N-degron pathway^45,46^.

### *In vivo* analysis of UBP6 cleavage in *Nicotiana benthamiana*

*Nicotiana benthamiana* leaves transiently expressing p35S::UBP6-GFP (pFASTR06-UBP6) using *Agrobacterium tumefaciens*-mediated infiltration were harvested (approximately 2–3 g), frozen in liquid nitrogen, and ground to a fine powder using a pre-chilled mortar and pestle. Protein extraction buffer (150 mM Tris-HCl pH 7.5, 150 mM NaCl, 10% glycerol, 10 mM EDTA, 10 mM EGTA, 50 μM Z-VRPR-fmk, 1 mM sodium molybdate, 1 mM NaF, 10 mM DTT, 0.5% (w/v) PVPP, 1× protease inhibitor cocktail [P9599, Sigma], and 1% (v/v) NP-40) was added at a ratio of 2 mL per gram of tissue. Samples were allowed to thaw on ice while being gently mixed until a homogeneous lysate was obtained. The lysate was clarified by centrifugation at 13,000 rcf for 20 min at 4°C, and the supernatant was filtered through a 40-µm cell strainer. For immunoprecipitation, 60 µL of GFP-Trap® Magnetic Beads (ChromoTek) slurry was thoroughly resuspended by gentle inversion and transferred to a LoBind protein1.5-mL tube using cut pipette tips. The beads were washed three times with 700 µL of wash buffer (20 mM Tris-HCl pH 7.5, 150 mM NaCl, 10 mM EDTA, 10 mM EGTA, 50 μM Z-VRPR-fmk and 0.5% [v/v] NP-40). For each wash, the beads were mixed by inversion, collected on a magnetic rack for 1-2 minutes and the supernatant was discarded. The washed beads were incubated with the clarified protein extract for 2 h at 4°C on a rotating wheel. Following incubation, the beads were collected on a magnetic rack and washed three times with 1.5 mL of wash buffer to remove unbound proteins and split in equal amounts for further analysis. The beads were then resuspended and washed in metacaspase protease reaction buffer (50 mM HEPES pH 7.5, 150 mM NaCl, 10% (w/v) glycerol, 50 mM CaCl2, and 10 mM DTT) for three times. Following immunoprecipitation, on-bead proteolysis was performed by incubating the GFP-Trap beads with the addition of either active or catalytically inactive recombinant MC9 protein produced in *Escherichia coli*, as previously described^28^. Proteolytic reactions were terminated by adding Laemmli sample buffer, and cleavage products were subsequently analysed by SDS–PAGE followed by immunoblotting.

### Pathogen inoculation in plant material

Bacterial suspension preparation and inoculation by injection or spraying were carried out on leaves of 3.5-week-old plants as described previously^13^. For gene expression and Western blot assays *Pst* DC3000, *Pst* DC3000 *avrRpm1*, and *Pst* DC3000 *hrcC−* were injected with a bacterial suspension of 10^7^ cfu/ml (OD_600 nm_ 0.1 = 10^8^ cfu/ml). For analysis of bacterial growth *Pst* DC3000 was injected or sprayed with bacterial suspensions of 10^6^ cfu/ml and 10^8^ cfu/ml, respectively. Three leaves per plant of at least seven plants were analysed.

### Extraction of proteins and Western blotting

Protein extractions and immunoblotting were carried out as described previously^13^. Primary antibodies used were mouse anti-HA (Sigma-Aldrich; 1:2,000), anti-NPR1 (donated by Dr. Steven Spoel; 1:500), anti-Ubiquitin (Santa Cruz Biotechnology; 1:1000), anti-RPN6 (Agrisera, 1:1000) and anti-Phospho-p44/42 MAPK (Erk1/2) (Thr202/Tyr204) (Cell Signaling Technology; 1:3,000). Horseradish peroxidase conjugated secondary antibodies, goat anti-Rabbit IgG (Agrisera) and goat anti-mouse IgG1 (Thermo Fisher Scientific), were used at 1:10,000 dilution. Signals were detected using Pierce™ ECL Western Blotting Substrate (Thermo Fisher Scientific) and imaged with a ChemiDoc MP Imaging System (Bio-Rad). Membranes were subsequently stained with Ponceau S, washed, dried, and scanned using a Epson Perfection V700 Photo scanner.

### Yeast two hybrid assay

E157-UBP6 and RPN6 cDNA sequences (without the STOP codon) were amplified from 7-day-old seedling cDNA and recombined into pDONR207. The constructs were mobilized into pGBKT7 vector (Gal4 DNA binding domain, BD; Clontech) and pGADT7 vector (Gal4 activation domain, AD; Clontech). The interaction of E157-UBP6 with RPN6 was assessed in vitro using the yeast two hybrid technique as previously described^47^.

### Coimmunoprecipitation assays

UBP6 *p6u6u7*, E157-UBP6 *p6u6u7* and *p6u6u7* seedlings were treated *in vitro* with SA 0.5mM for 24h. Coimmunoprecipitation assays were carried out as described previously with modifications^47^. Normalized seedling protein extracts were incubated with 25 μL of HA-Trap® magnetic beads (Chromotek) for 3 h at 4 °C with rotation. After washing five times with 500 μL of extraction buffer, samples were denatured by boiling in Laemmli sample buffer, separated on SDS-PAGE gels, and transferred onto polyvinylidene difluoride membranes (PVDF) (Bio-Rad).

### Affinity purification-Mass spectrometry

UBP6 *prt6-1*, E157-UBP6 *prt6-1* and *prt6-1* seedlings were treated in vitro with SA 0.5mM for 24h. Immunoprecipitation was performed as indicated above (Coimmunoprecipitation assays).

#### S-TrapTM Digestion

Protein concentration was estimated by Pierce 660nm protein assay (Thermo Fisher Scientific). Protein digestion in the S-Trap filter (Protifi, Huntington, NY, USA) was performed following the protocol described in^48^ with slight modifications. Briefly, 20 µg of protein of each sample was diluted to 40 µL with 5% SDS. Samples were reduced and alkylated by adding 5 mM tris(2-carboxyethyl)phosphine and 10 mM chloroacetamide for 30 minutes at 60°C. Afterwards, 12% phosphoric acid and then seven volumes of binding buffer (90% methanol; 100 mM TEAB) were added to the sample (final phosphoric acid concentration: 1.2%). After mixing, the protein solution was loaded to an S-Trap filter in two consecutive steps, separated by a 2 min centrifugation at 3000 x g. Then the filter was washed 3 times with 150 μL of binding buffer. Finally, samples were digested at 37°C with Pierce MS-grade trypsin (Thermo-Fisher Scientific) at an enzyme:substrate ratio of 1:20 (w/w) in 20 μL of a 100 mM TEAB solution. Samples were spun through the S-Trap prior to digestion. Flow-through was then reloaded to the top of the S-Trap column and allowed to digest in a wet chamber at 37°C overnight. To elute peptides, two step-wise buffers were applied (1) 40 μL of 25 mM TEAB and 2) 40 μL of 80% acetonitrile and 0.2% formic acid in H2O), separated by a 2 min centrifugation at 3000 x g in each case. Eluted peptides were pooled and vacuum centrifuged to dryness.

#### Liquid chromatography and mass spectrometry analysis

*(LC-ESI-MS/MS)*: Digested samples were cleaned-up/desalted using Stage-Tips with Empore 3M C18 disks (Sigma-Aldrich) [1]. After desalting, peptide concentration was carried out by Qubit™ Fluorometric Quantitation (Thermo Fisher Scientific). A 500ng aliquot of each digested sample was subjected to 1D-nano LC ESI-MS/MS (Liquid Chromatography Electrospray Ionization Tandem Mass Spectrometric) analysis using an Ultimate 3000 nano HPLC system (Thermo Fisher Scientific) coupled online to a Orbitrap Exploris 240 equipped with FAIMS Pro ion source (Thermo Fisher Scientific). Peptides were eluted onto a 50 cm × 75 μm Easy-spray PepMap C18 analytical column at 45°C and were separated at a flow rate of 250 nL/min using a 60 min gradient ranging from 2 % to 95 % mobile phase B (mobile phase A: 0.1% formic acid (FA); mobile phase B: 100 % acetonitrile (ACN), 0.1 % FA). The loading solvent was 2 % ACN) in 0.1 % FA and injection volume was 5 µl.

Data acquisition was performed using a data-dependent top-20 method, in full scan positive mode, scanning 375 to 1200 m/z. Survey scans were acquired at a resolution of 60,000 at m/z 200, with Normalized Automatic Gain Control (AGC) target (%) of 300 and a maximum injection time (IT) of 40 ms. The top 20 most intense ions from each MS1 scan were selected and fragmented via Higher-energy collisional dissociation (HCD). Resolution for HCD spectra was set to 45,000 at m/z 200, with AGC target of 200 and maximum ion injection time of 120 ms. Isolation of precursors was performed with a window of 1 m/z, exclusion duration (s) of 45 and the HCD collision energy was 32. Precursor ions with single, unassigned, or six and higher charge states from fragmentation selection were excluded.

*Proteomics data analysis and sequence search*: Raw instrument files were processed using Proteome Discoverer (PD) version 3.1 (Thermo Fisher Scientific). MS2 spectra were searched using Mascot Server v3.1 (Matrix Science, London, UK) and a target/decoy database built from sequences in the Arabidopsis thaliana (UKBsp_p3702Ref). All searches were configured with dynamic modifications for pyrrolidone from Q (−17.027 Da) and oxidation of methionine residues (+15.9949 Da) and static modification as carbamidomethyl (+57.021 Da) on cysteine, monoisotopic masses, and trypsin cleavage (max 2 missed cleavages). The peptide precursor mass tolerance was 10 ppm, and MS/MS tolerance was 0.02 Da. The false discovery rate (FDR) for proteins, peptides, and peptide spectral matches (PSMs) peptides were kept at 1%.

### Gene expression analyses

Total RNA was extracted from seedlings and plants (age indicated for each experiment) using a Plant Total RNA Mini Kit (FAVORPREP) following the manufactureŕs instructions. cDNA synthesis and quantitative RT-PCR were performed as previously described for Arabidopsis ^49^ using a High-Capacity cDNA Reverse Transcription Kit (Thermo Fisher Scientific) and qPCR Hot Firepol Evagreen qPCR Mix Plus 5x (Cultek). For primers used see Supporting Information Table 1.

### SA RNA-Seq

Total RNA from seedlings treated *in vitro* with SA 0.5mM was extracted, as indicated above. Four independent assays, 40 seedlings per line and treatment in each assay. Mock samples were subjected to the same process as the SA-treated. Briefly, mRNA was isolated with poly-T oligo-attached magnetic beads and fragmented. First strand cDNA was synthesized using random hexamer primers followed by second strand cDNA synthesis. Library quality was checked with a Qubit fluorometer (Thermo Fisher) and quantified by real-time PCR. Quantified libraries were pooled and paired-end sequenced on the Illumina platform NovaSeq X Plus (PE150). Raw reads were filtered to remove adapters and low-quality reads. In all samples, over 95% of reads had an error rate of less than 0.001. Filtered reads were mapped to the Arabidopsis thaliana genome (TAIR10 release) using hisat2^50^. Alignments were visually inspected with the IGV browser^51^. Transcripts were quantified with featureCounts^52^ (parameters: -s 0 -p –countReadPairs -F SAF -g gene -a Arabidopsis_thaliana.TAIR10.57.saf). For each comparison, differentially expressed genes were detected with the DESeq2 Bioconductor package^53^ using default options.

### Phylogenetic analysis

Phylogenetic analyses of UBP6-like sequences were performed by identifying orthologous proteins in different plant and animal species using BLASTP on the NCBI server. The retrieved sequences were aligned using Clustal Omega, the resulting alignments were visualized with Jalview v2.11.5.1, and the phylogenetic tree was subsequently constructed using FastTree on the Phylogeny.fr server. In addition, sequence logos were generated from the regions of interest using the Leskoff Sequence Logo Generator.

### Production of recombinant MC9 protein

MC9 open reading frame was cloned into pDEST17 vector from *A. thaliana* seedling RNA, resulting in an N-terminal fusion with a His x 6 epitope tag. Protein was induced at 30 °C overnight with 1 mM IPTG and purified as described previously^21^.

### MC9 activity assay *in vitro*

We investigated the direct proteolytic effect of MC9 on UBP6 by performing an *in vitro* cleavage assay using recombinant MC9 and UBP6 substrates produced by coupled transcription/translation. UBP6 WT and UBP6(R156A) in the vector UFT-3HA were used as templates for *in vitro* transcription and translation (TnT® T7/T3 Coupled Reticulocyte Lysate System, Promega) according to the manufacturer’s instructions. Reactions were gently mixed on ice and incubated at 30 °C for 30 min. After incubation, 2 μl of 2.6 mM cycloheximide (Sigma-Aldrich) was added to inhibit translation. Each reaction was divided into two 15 μl aliquots, one for the MC9 treatment (+MC9) and one as a negative control (−MC9). Recombinant MC9 was pre-activated prior to use in activity buffer (50 mM MES, pH 5.5, 150 mM NaCl, 10% (w/v) sucrose, 0.1% (w/v) CHAPS)^54^. For pre-activation, 20 μl activity buffer and 10 μl 100 mM DTT were added to 50 μl of recombinant MC9 (2.938 mg/mL, ∼ 63.9 µM), and the mixture was incubated at 30 °C for 10 min. For cleavage assays, 30 μl of pre-activated MC9 was added to 15 μl of the in vitro–translation solution, followed by 10 μl activity buffer. A 25 μl aliquot was collected immediately (time 0 min). Negative control reactions contained 40 μl activity buffer in place of MC9. Reactions were incubated at 30 °C, and additional samples were subsequently collected. Reaction products were denatured by addition of 7 μl Laemmli sample buffer and water to a final volume of 42 μl, followed by incubation at 95 °C for 10 min, chilling on ice for 5 min, brief centrifugation, and storage at −20 °C.

### Structure prediction and simulations

We built full-sequence models of AtUBP6 using C-I-TASSER^55^, which produced a fully folded structure that closely matched known ScUbp6 structures in PDB entries 7QO3 and 7QO4^56^. Deletion mutants E157 and S141 were generated by homology modelling using the model for UBP6 as reference with Modeller^57^ and UCSF Chimera^58^, with thorough optimization to produce 5 models with hydrogens, and selecting the best. To further refine the structures and investigate their potential stability, they were subjected to a Molecular Dynamics (MD) simulation using GROMACS^59^ at pH 7.365, 1 atm, 310°K in saline solution for 10 ns with the Amber force field^60^. The simulations were inspected and representative structures selected for modelling Ubp6 bound to the proteasome: refined MD models were used to substitute UBP6 in PDB structure 7QO4 which corresponds to the 26S proteasome WT-Ubp6-UbVS complex in the si state (*S. cerevisiae*, Human). The structures were then subjected to an initial short minimization using UCSF Chimera and AMBER followed by longer minimizations in GROMACS with the AMBER force field first in vacuo and then in saline solution to allow them to adapt mutually before inspection.

### Experimental statistical analyses

All experiments were performed at least in triplicate. Statistical comparisons were conducted using GraphPad Prism 9.0 software and R. Horizontal lines represent standard error of the mean values in all graphs. For two-sample comparisons we used Student’s *t*-test or Wilcoxon test, where statistically significant differences are reported as: ***, *P* < 0.001; **, *P* < 0.01; *, *P* < 0.05; For multiple comparisons we used and one-way analysis of variance (ANOVA) and Tukey’s post-hoc test, where significant differences (α < 0.05) are denoted with different letters.

## Data availability

RNA-seq data have been deposited in the NCBI Gene Expression Omnibus (GEO) under accession number GSE335550. The data are currently private and available to reviewers using the following access token: ubsdocgyzzmzdin.

Data and resources will to be shared upon reasonable request to Jorge Vicente and Michael Holdsworth.

## Acknowledgements

We thank Beatriz Casal, María Luisa Peinado and Daniel García for technical support of this work. The authors acknowledge the Proteomics Facility of the Spanish National Center for Biotechnology (CNB-CSIC) for technical support and for performing the LC-ESI-MS/MS proteomic analyses.

## Author information

### Contributions

J.V. and M.J.H conceived the research with inputs from C.D, G.S., FvB, K.G, N.O. All authors contributed towards design of experiments and interpretation of data; J.V., M.J.H, C.D, G.S., I.M, Y.I., A.D.F, J.R.V. and N.J.O. carried out the research; J.V. and M.J.H wrote the manuscript. J.V. agrees to serve as the author responsible for contact and ensures communication. All authors read, contributed to editing and approved the manuscript.

## List of Extended Data

Supplemental Figure 1: Current knowledge of the role of the N-Degron pathways in plant biotic stress responses and representation of the protein domain within the Ubiquitin Fusion Technique (UFT) construct used in this work.

Supplemental Figure 2: Data related to impairment of the N-degron pathways enhances plant response to PAMPs, *MC9* expression and conservation and structure in the region spanning A139–R156.

Supplemental Figure 3: E157-UBP6 interacts with the proteasome and inhibits its activity. Supplemental Data 1: List of proteins that interacts with E157-UBP6 and full-length UBP6 after SA treatment.

Supplemental Data 2: DEGs in E157-UBP6 *p6u6u7* compared with *p6u6u7* after SA treatment.

Supplemental Table 1: List of oligonucleotide primer sequences used for cloning, qRT-PCR and genotyping.

