## Supplemental Figures for "Functional diversification of UBP6 in plant immunity through N-degron pathway regulation"

### Slide 1
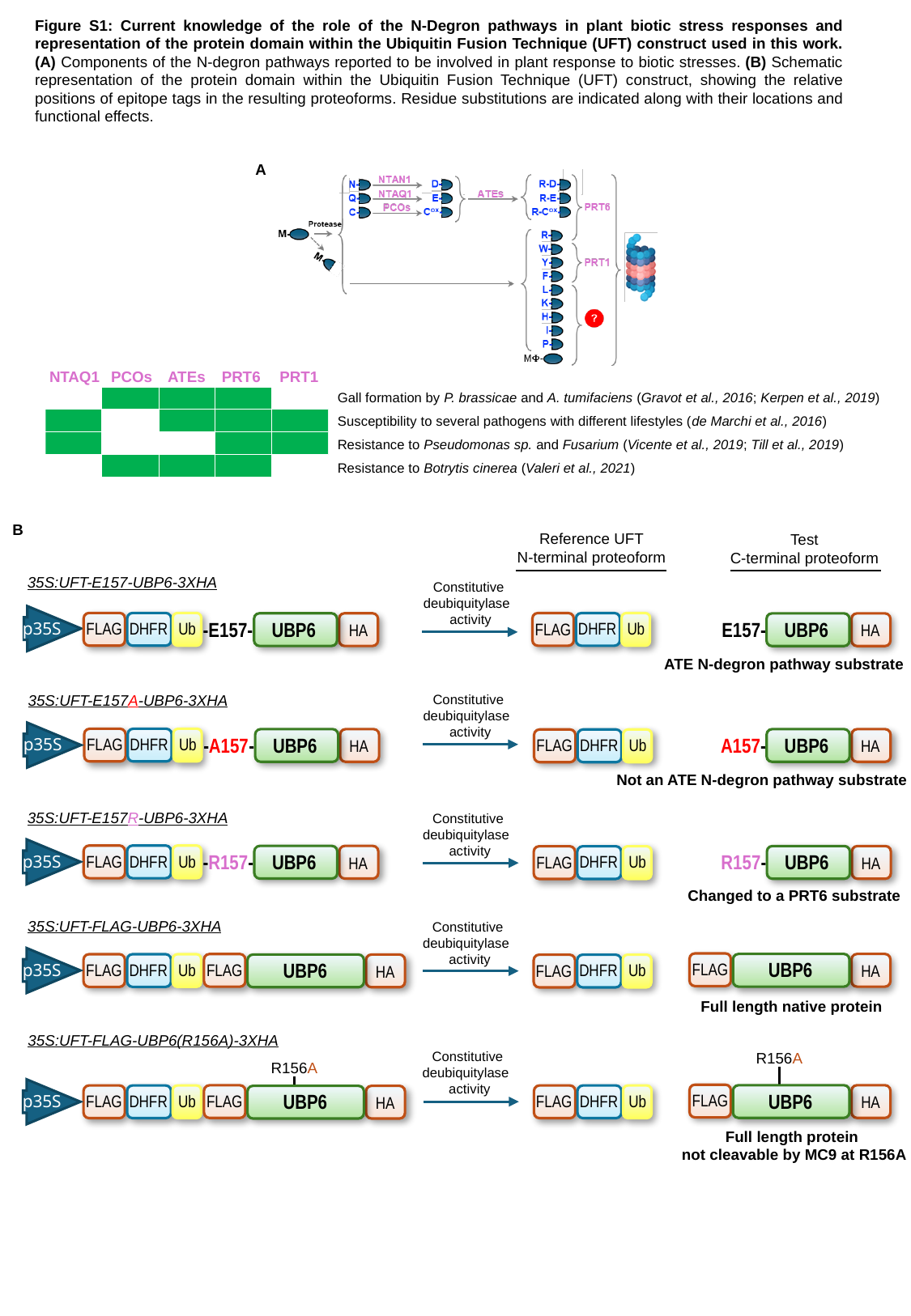

Figure S1: Current knowledge of the role of the N-Degron pathways in plant biotic stress responses and representation of the protein domain within the Ubiquitin Fusion Technique (UFT) construct used in this work. (A) Components of the N-degron pathways reported to be involved in plant response to biotic stresses. (B) Schematic representation of the protein domain within the Ubiquitin Fusion Technique (UFT) construct, showing the relative positions of epitope tags in the resulting proteoforms. Residue substitutions are indicated along with their locations and functional effects.
A
NTAQ1
PCOs
ATEs
PRT6
PRT1
Gall formation by P. brassicae and A. tumifaciens (Gravot et al., 2016; Kerpen et al., 2019)
Susceptibility to several pathogens with different lifestyles (de Marchi et al., 2016)
Resistance to Pseudomonas sp. and Fusarium (Vicente et al., 2019; Till et al., 2019)
Resistance to Botrytis cinerea (Valeri et al., 2021)
B
Reference UFT
N-terminal proteoform
Test
C-terminal proteoform
35S:UFT-E157-UBP6-3XHA
Constitutive
deubiquitylase
activity
-E157-
UBP6
E157-
UBP6
Ub
DHFR
p35S
Ub
DHFR
FLAG
FLAG
HA
HA
ATE N-degron pathway substrate
35S:UFT-E157A-UBP6-3XHA
Constitutive
deubiquitylase
activity
-A157-
A157-
UBP6
UBP6
Ub
DHFR
p35S
FLAG
Ub
DHFR
FLAG
HA
HA
Not an ATE N-degron pathway substrate
35S:UFT-E157R-UBP6-3XHA
Constitutive
deubiquitylase
activity
-R157-
UBP6
R157-
UBP6
Ub
DHFR
p35S
FLAG
Ub
DHFR
FLAG
HA
HA
Changed to a PRT6 substrate
35S:UFT-FLAG-UBP6-3XHA
Constitutive
deubiquitylase
activity
UBP6
UBP6
FLAG
Ub
FLAG
DHFR
p35S
FLAG
Ub
DHFR
FLAG
HA
HA
Full length native protein
35S:UFT-FLAG-UBP6(R156A)-3XHA
Constitutive
deubiquitylase
activity
R156A
R156A
UBP6
UBP6
FLAG
Ub
Ub
FLAG
DHFR
DHFR
p35S
FLAG
FLAG
HA
HA
Full length protein
not cleavable by MC9 at R156A

### Slide 2
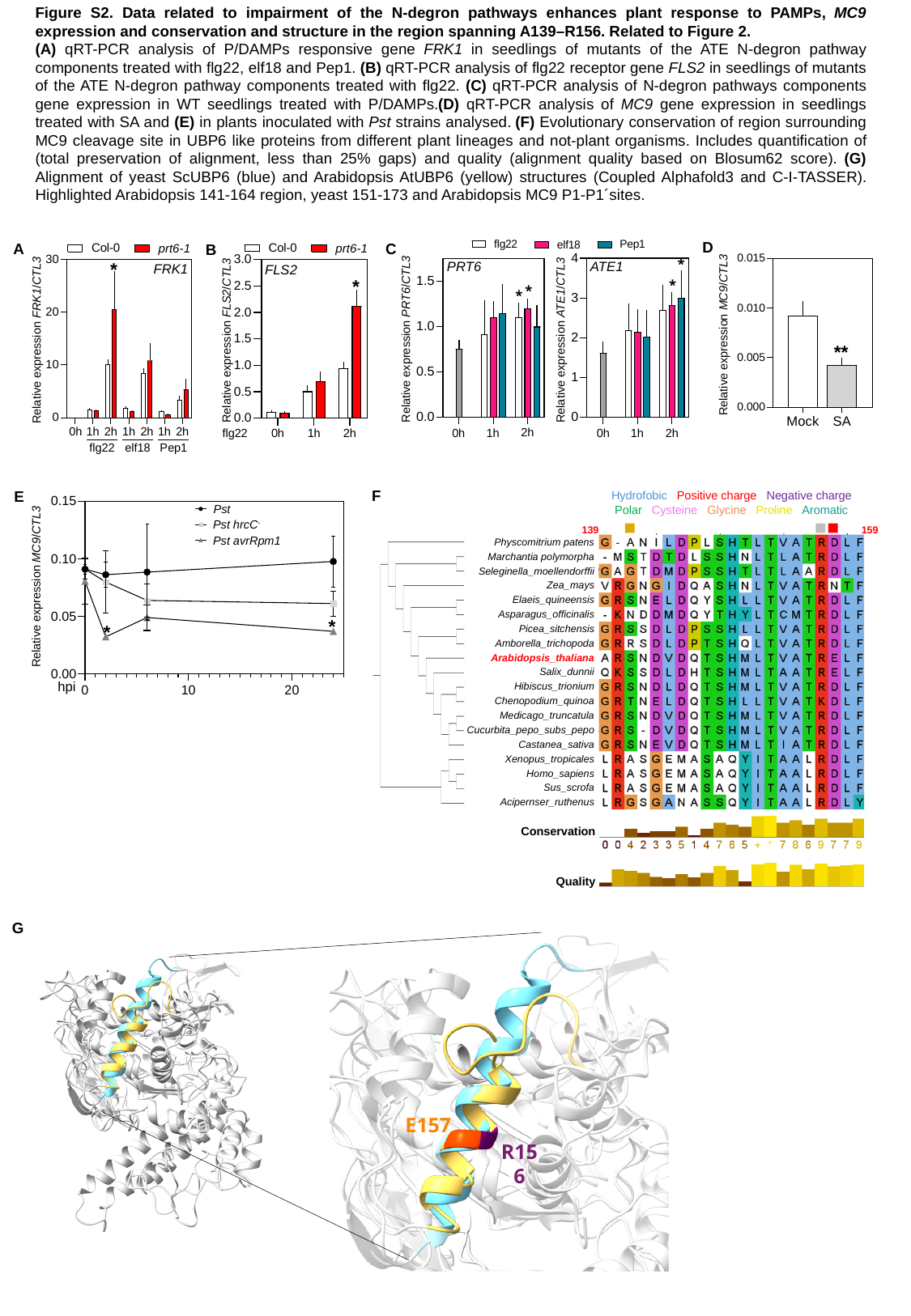

Figure S2. Data related to impairment of the N-degron pathways enhances plant response to PAMPs, MC9 expression and conservation and structure in the region spanning A139–R156. Related to Figure 2.
(A) qRT-PCR analysis of P/DAMPs responsive gene FRK1 in seedlings of mutants of the ATE N-degron pathway components treated with flg22, elf18 and Pep1. (B) qRT-PCR analysis of flg22 receptor gene FLS2 in seedlings of mutants of the ATE N-degron pathway components treated with flg22. (C) qRT-PCR analysis of N-degron pathways components gene expression in WT seedlings treated with P/DAMPs.(D) qRT-PCR analysis of MC9 gene expression in seedlings treated with SA and (E) in plants inoculated with Pst strains analysed. (F) Evolutionary conservation of region surrounding MC9 cleavage site in UBP6 like proteins from different plant lineages and not-plant organisms. Includes quantification of (total preservation of alignment, less than 25% gaps) and quality (alignment quality based on Blosum62 score). (G) Alignment of yeast ScUBP6 (blue) and Arabidopsis AtUBP6 (yellow) structures (Coupled Alphafold3 and C-I-TASSER). Highlighted Arabidopsis 141-164 region, yeast 151-173 and Arabidopsis MC9 P1-P1´sites.
D
C
A
B
*
*
PRT6
ATE1
FRK1
FLS2
*
*
*
*
Relative expression MC9/CTL3
Relative expression PRT6/CTL3
Relative expression ATE1/CTL3
Relative expression FRK1/CTL3
Relative expression FLS2/CTL3
**
Mock
SA
2h
2h
2h
1h
0h
1h
1h
2h
1h
0h
2h
1h
0h
flg22
2h
1h
0h
Pep1
flg22
elf18
F
E
Hydrofobic Positive charge Negative charge Polar Cysteine Glycine Proline Aromatic
Pst
Pst hrcC^-^
159
139
Pst avrRpm1
Physcomitrium patens
Marchantia polymorpha
Seleginella_moellendorffii
Relative expression MC9/CTL3
Zea_mays
Elaeis_quineensis
Asparagus_officinalis
*
*
Picea_sitchensis
Amborella_trichopoda
Arabidopsis_thaliana
Salix_dunnii
hpi
Hibiscus_trionium
Chenopodium_quinoa
Medicago_truncatula
Cucurbita_pepo_subs_pepo
Castanea_sativa
Xenopus_tropicales
Homo_sapiens
Sus_scrofa
Acipernser_ruthenus
Conservation
Quality
G
E157
R156

### Slide 3
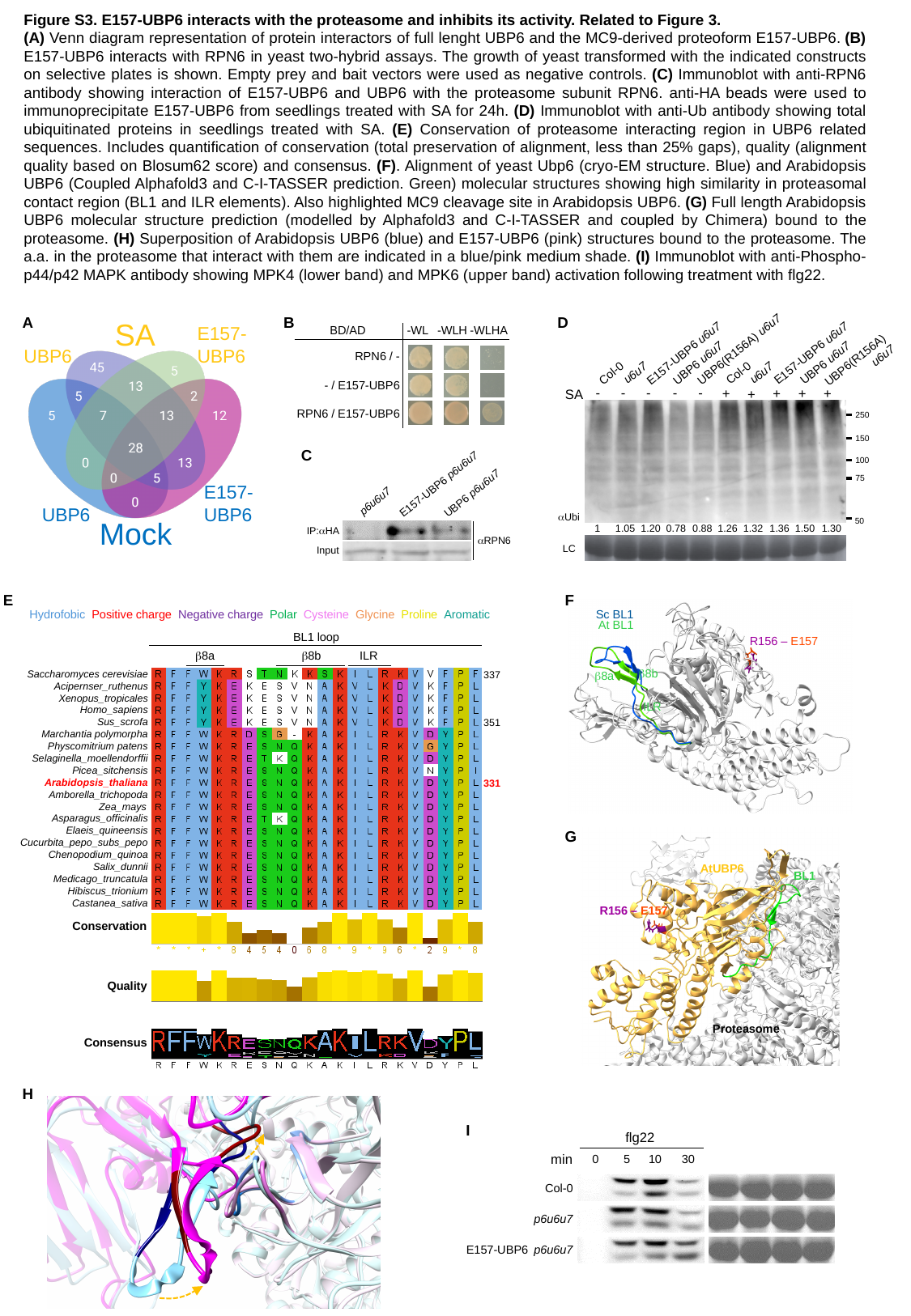

Figure S3. E157-UBP6 interacts with the proteasome and inhibits its activity. Related to Figure 3.
(A) Venn diagram representation of protein interactors of full lenght UBP6 and the MC9-derived proteoform E157-UBP6. (B) E157-UBP6 interacts with RPN6 in yeast two-hybrid assays. The growth of yeast transformed with the indicated constructs on selective plates is shown. Empty prey and bait vectors were used as negative controls. (C) Immunoblot with anti-RPN6 antibody showing interaction of E157-UBP6 and UBP6 with the proteasome subunit RPN6. anti-HA beads were used to immunoprecipitate E157-UBP6 from seedlings treated with SA for 24h. (D) Immunoblot with anti-Ub antibody showing total ubiquitinated proteins in seedlings treated with SA. (E) Conservation of proteasome interacting region in UBP6 related sequences. Includes quantification of conservation (total preservation of alignment, less than 25% gaps), quality (alignment quality based on Blosum62 score) and consensus. (F). Alignment of yeast Ubp6 (cryo-EM structure. Blue) and Arabidopsis UBP6 (Coupled Alphafold3 and C-I-TASSER prediction. Green) molecular structures showing high similarity in proteasomal contact region (BL1 and ILR elements). Also highlighted MC9 cleavage site in Arabidopsis UBP6. (G) Full length Arabidopsis UBP6 molecular structure prediction (modelled by Alphafold3 and C-I-TASSER and coupled by Chimera) bound to the proteasome. (H) Superposition of Arabidopsis UBP6 (blue) and E157-UBP6 (pink) structures bound to the proteasome. The a.a. in the proteasome that interact with them are indicated in a blue/pink medium shade. (I) Immunoblot with anti-Phospho-p44/p42 MAPK antibody showing MPK4 (lower band) and MPK6 (upper band) activation following treatment with flg22.
A
B
D
SA
E157-
UBP6
-WL
-WLH
-WLHA
BD/AD
UBP6(R156A) u6u7
E157-UBP6 u6u7
E157-UBP6 u6u7
UBP6
RPN6 / -
UBP6(R156A)
u6u7
UBP6 u6u7
UBP6 u6u7
Col-0
Col-0
u6u7
u6u7
- / E157-UBP6
-
-
-
-
-
+
+
+
+
+
SA
RPN6 / E157-UBP6
250
150
C
100
75
E157-UBP6 p6u6u7
E157-
UBP6
UBP6 p6u6u7
p6u6u7
UBP6
αUbi
Mock
50
1
1.05
1.20
0.78
0.88
1.26
1.32
1.36
1.50
1.30
IP:αHA
αRPN6
LC
Input
E
F
Hydrofobic Positive charge Negative charge Polar Cysteine Glycine Proline Aromatic
Sc BL1
At BL1
BL1 loop
R156 – E157
β8a
ILR
β8b
β8b
Saccharomyces cerevisiae
337
β8a
Acipernser_ruthenus
Xenopus_tropicales
ILR
Homo_sapiens
Sus_scrofa
351
Marchantia polymorpha
Physcomitrium patens
Selaginella_moellendorffii
Picea_sitchensis
Arabidopsis_thaliana
331
Amborella_trichopoda
Zea_mays
Asparagus_officinalis
Elaeis_quineensis
G
AtUBP6
BL1
R156 – E157
Proteasome
Cucurbita_pepo_subs_pepo
Chenopodium_quinoa
Salix_dunnii
Medicago_truncatula
Hibiscus_trionium
Castanea_sativa
Conservation
Quality
Consensus
H
I
flg22
min
0
5
10
30
Col-0
p6u6u7
E157-UBP6
p6u6u7
